# Genome-scale phylogenetic analyses confirm *Olpidium* as the closest living zoosporic fungus to the non-flagellated, terrestrial fungi

**DOI:** 10.1101/2020.09.16.298935

**Authors:** Ying Chang, D’Ann Rochon, Satoshi Sekimoto, Yan Wang, Mansi Chovatia, Laura Sandor, Asaf Salamov, Igor V. Grigoriev, Jason E. Stajich, Joseph W. Spatafora

## Abstract

The zoosporic obligate endoparasites, *Olpidium*, hold a pivotal position to the reconstruction of the flagellum loss in fungi, one of the key morphological transitions associated with the colonization of land by the early fungi. We generated genome and transcriptome data from non-axenic zoospores of *Olpidium bornovanus* and used a metagenome approach to extract phylogenetically informative fungal markers. Our phylogenetic reconstruction strongly supported *Olpidium* as the closest zoosporic relative of the non-flagellated terrestrial fungi. Super-alignment analyses resolved *Olpidium* as sister to the non-flagellated terrestrial fungi, whereas a super-tree approach recovered different placements of *Olpidium*, but without strong support. Further investigations detected little conflicting signal among the sampled markers but revealed a potential polytomy in early fungal evolution associated with the branching order among *Olpidium*, Zoopagomycota and Mucoromycota. The branches defining the evolutionary relationships of these lineages were characterized by short branch lengths and low phylogenetic content and received equivocal support for alternative phylogenetic hypotheses from individual markers. These nodes were marked by important morphological innovations, including the transition to hyphal growth and the loss of flagellum, which enabled early fungi to explore new niches and resulted in rapid and temporally concurrent Precambrian diversifications of the ancestors of several phyla of fungi.

## Introduction

Accuracy of the higher-level relationships within Kingdom Fungi has improved with the analysis of increasingly more samples of whole genome sequence data. Several problematic lineages have been placed with confidence in the tree of fungi, including the grouping of Microsporidia with Cryptomycota ^1^, the inclusion of arbuscular mycorrhizal fungi (Glomeromycotina) in Mucoromycota ^2^, and the placement of the xerophilic Wallemiomycetes as an early branch in Agaricomycotina ^3^. On the other hand, some deep nodes in the fungal phylogeny remain unresolved despite the recent efforts on phylogenomic studies. The relationships among the three subphyla within Basidiomycota could not be resolved with dense sampling of whole genome data ^4^, and while genome scale data supported the delineation of Mucoromycota and Zoopagomycota, the branching order of their respective subphyla remains unclear. Similarly, genome-scale phylogeny supports the recognition of separate phyla of zoosporic fungi Blastocladiomycota and Chytridiomycota, but resolution of their branching order remains elusive ^5^.

Resolving deep divergence events has been challenging for a number of reasons. Due to the ancient ages of these events, the evolutionary signals have eroded through time with multiple substitutions accumulated at the same sites ^6^. When rapid radiation events were associated with these deep nodes, their reconstruction is even more difficult because less information can be mapped onto the short internodes and the inference of these nodes are also more prone to stochastic errors ^7^. Another source of lack of resolution is the inconsistent signals contained in individual markers. These inconsistent signals could come from real biological sources, such as ancient hybridization events, horizontal gene transfers, introgressions, or incomplete lineage sorting ^8,9^. Even when individual markers share the same evolutionary history, the inferred phylogeny of the genes can still differ due to systematic bias introduced by improper modeling of data or analytical artifact of long branch attraction (LBA) caused by presence of extremely long terminal branches ^6^. One effective way to reduce the systematic errors and improve phylogenetic estimations is to increase taxonomic sampling ^10,11^. Dense taxon sampling can help break down long terminal branches and therefore can mitigate the effect of LBA. Increased taxonomic sampling can also help optimize model parameter estimates and improve phylogenetic accuracy ^12,13^.

The loss of the flagellum and the origin of a hyphal body plan are among the most important morphological transitions in fungal evolution ^14^, and the genus *Olpidium* is critical in the understanding of flagellum loss in fungi. Despite being zoosporic, *Olpidium* was placed within the non-flagellated terrestrial fungi by multi-gene phylogenetic reconstructions ^15,16^, albeit without strong support. The posterior flagellum of the zoospores of fungi is a synapomorphy shared by fungi and other the opisthokont relatives. Liu et al. ^17^ hypothesized that the flagellum was lost just once in the evolution of terrestrial fungi, and that this coincided with shift of microtubular organization center (MTOC) from centrioles to spindle pole bodies (SPBs). The placement of *Olpidium* with fungi of Zoopagomycota in James *et al*.^18^ and Sekimoto *et al*. ^16^, on the other hand, suggests that there were more than one flagellum loss events. Therefore, accurate placement of *Olpidium* in the fungal phylogeny will not only help us to estimate the number of flagellum losses in the evolution of fungi, but also improve the understanding of MTOC evolution among the early-diverging fungal lineages.

*Olpidium* includes about 30 described species, many of them obligate endoparasites of algae, plants, fungi and small animals. Morphologically *Olpidium* resembles the typical ‘core chytrid’, with a holocarpic thallus that produces zoospores with a single posterior flagellum ^19^. There has not been any genus-wide systematic study on *Olpidium*, and our understanding of *Olpidium* biology has largely come from pathogenicity and ecological studies on three species, *O. bornovanus, O. brassicae* and *O. virulentus* ^20–23^, which are root parasites of higher plants, including many crops like cabbages and lettuce ^24^. The infection by these *Olpidium* species generally does not cause any symptoms to the host plants. However, the zoospores of *O. bornovanus* and *O. virulentus* can function as vectors of viruses (e.g., Mirafiori lettuce big-vein virus), which often cause considerable damages to the host plants ^25^. These *Olpidium* species can sometimes make up a large proportion of the fungal communities associated with plant roots or the rhizosphere ^26^.

Collecting sufficient amounts of tissue for DNA extraction remains one of the major challenges for phylogenomic studies on obligate fungal symbionts. For species that cannot be maintained in axenic culture, DNA data can only be generated from field collected specimens like fruiting bodies or from fungal tissue co-cultured with the hosts. Here, we collected DNA from zoospores of *Olpidium bornovanus* that were produced in bulk after repeated rounds of inoculation of cucumber roots followed by induction of zoospore release. The DNA data from obligate symbionts is often from heterogeneous origins and requires careful data binning and accurate taxonomic assignment before any subsequent analysis. While the majority of the metagenomic tools are designed for binning bacterial genomes, studies have also successfully applied metagenomic approaches to eukaryotic genomes ^27–29^. In this study, we performed whole-genome sequencing on *Olpidium* DNA derived from zoospore material collected from co-cultured plant roots. We employed a metagenomic approach and extracted sequence data from *Olpidium*. We retrieved 295 markers from *Olpidium* for phylogenetic reconstruction, together with dense taxon sampling in the zoosporic lineages, i.e., Chytridiomycota and Blastocladiomycota, and the early-diverging non-flagellated terrestrial fungi, i.e., Zoopagomycota and Mucoromycota (Table S1). In this study we addressed the placement of *Olpidium* in Kingdom Fungi with a phylogenomic approach. We also explored whether the recalcitrant nodes of early fungal evolution were caused mainly by the presence of insufficient signal or by incongruent phylogenetic information contained in individual genes.

## Results

### Genome assembly and data binning analysis

The raw *de novo* assembly of transcriptome comprised 94,984 contigs with a total length of 41 megabases (Mb). A large proportion of contigs from this assembly (80%) were less than 1.5 kb in length and not included in the binning analysis using VIZBIN ^30^. The subsampled transcriptome assembly from VIZBIN analysis, cluster 1 (Fig. 1), contained 1,946 sequences with a total of 4.4 Mb. The raw *de novo* assembly of *Olpidium bornovanus* genome comprised 160,327 contigs with a total length of 187 Mb. After two rounds of binning and conservative sampling of data (Fig. 1), the working genome assembly contained 13,695 contigs with a total length of 29 Mb.

**Figure 1.**
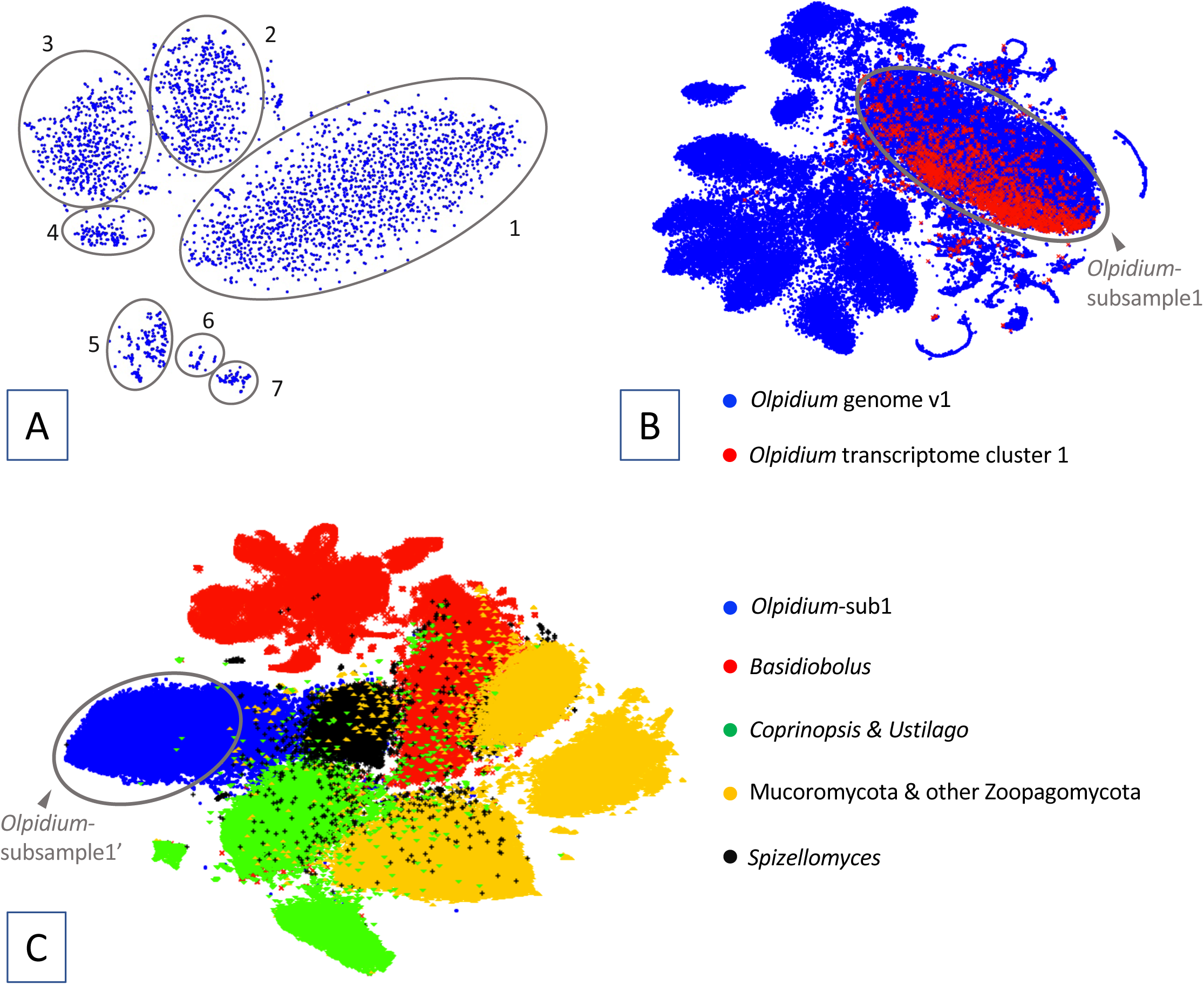
Metagenomic data binning of the transcriptome and genome assemblies of *Olpidium bornovanus* based on tetramer nucleotide composition using Vizbin. 1A: Vizbin analysis on the transcriptome. Contigs longer shorter than 1.5 kb were removed from the analysis. The ovals delimit the seven data clusters recovered in the Vizbin analysis. 1B: Vizbin analysis on the *Olpidium-*raw-genome assembly. Contigs longer than 1kb in the genome assembly (blue data points) and contigs from transcriptome cluster 1 (red data points) were included in the analysis. The oval delimits *Olpidium-*subsample1, the subset of contigs used in the third Vizbin analysis. 1C: Vizbin analysis on *Olpidium-*subsample1 (blue data points). Genome sequences from another seven fungal species were included in the same Vizbin analysis to serve as spike sequences. Contigs from the area that the *Olpidium*-subsample1 contigs show little overlap with the spike sequences were sampled (*Olpidium-*subsample1’) and further binned using methods based on sequence similarity (mmseq2 and DIAMOND blastx) to produce the working assembly.

### Phylogenetic reconstruction and dating analysis

The concatenated matrix of the 295-marker dataset comprised 83,461 aligned amino acid sites. The results based on RAxML ^31^ and IQ-TREE ^32^analyses were generally consistent with strong support for the monophyly of fungal phyla and subphyla sampled in this study (bootstrap support 100; Fig. 2 & Table S2; referred as ConcatBestTree). The first split in Kingdom Fungi was inferred between Cryptomycota and the remaining fungal taxa. Blastocladiomycota and Chytridiomycota formed a paraphyletic grade of zoosporic fungi, subtending a clade of *Olpidium* and the non-flagellated terrestrial lineages. All the splits mentioned above, as well as the *Olpidium* + non-flagellated clade, received strong support (bootstrap support of 100) from RAxML and IQ-TREE analyses. *Olpidium* was resolved as the sister group of non-flagellated terrestrial fungi with moderate to strong bootstrap support (95 for IQ-TREE analysis and 89 for RAxML analysis on non-partitioned 295-marker dataset; Fig. 2 & Table S2). Within non-flagellated terrestrial fungi, Zoopagomycota was sister to a clade of Mucoromycota and Dikarya, again with moderate to strong bootstrap support (95 for IQ-TREE analysis and 89 for RAxML on non-partitioned 295-marker dataset). Partitioned RAxML analysis on the 295-marker dataset showed weaker support for the aforementioned nodes (66 and 68, respectively). ASTRAL^33^ analyses also recovered the clade of *Olpidium* and the non-flagellated fungi with strong support (1.0 pp and 100 bootstrap support). For the exact placement of *Olpidium*, ASTRAL analyses showed decreased support for *Olpidium* as sister to non-flagellated group, and various placements of *Olpidium* were recovered in the best trees across different ASTRAL analyses (Fig. S1).

**Figure 2.**
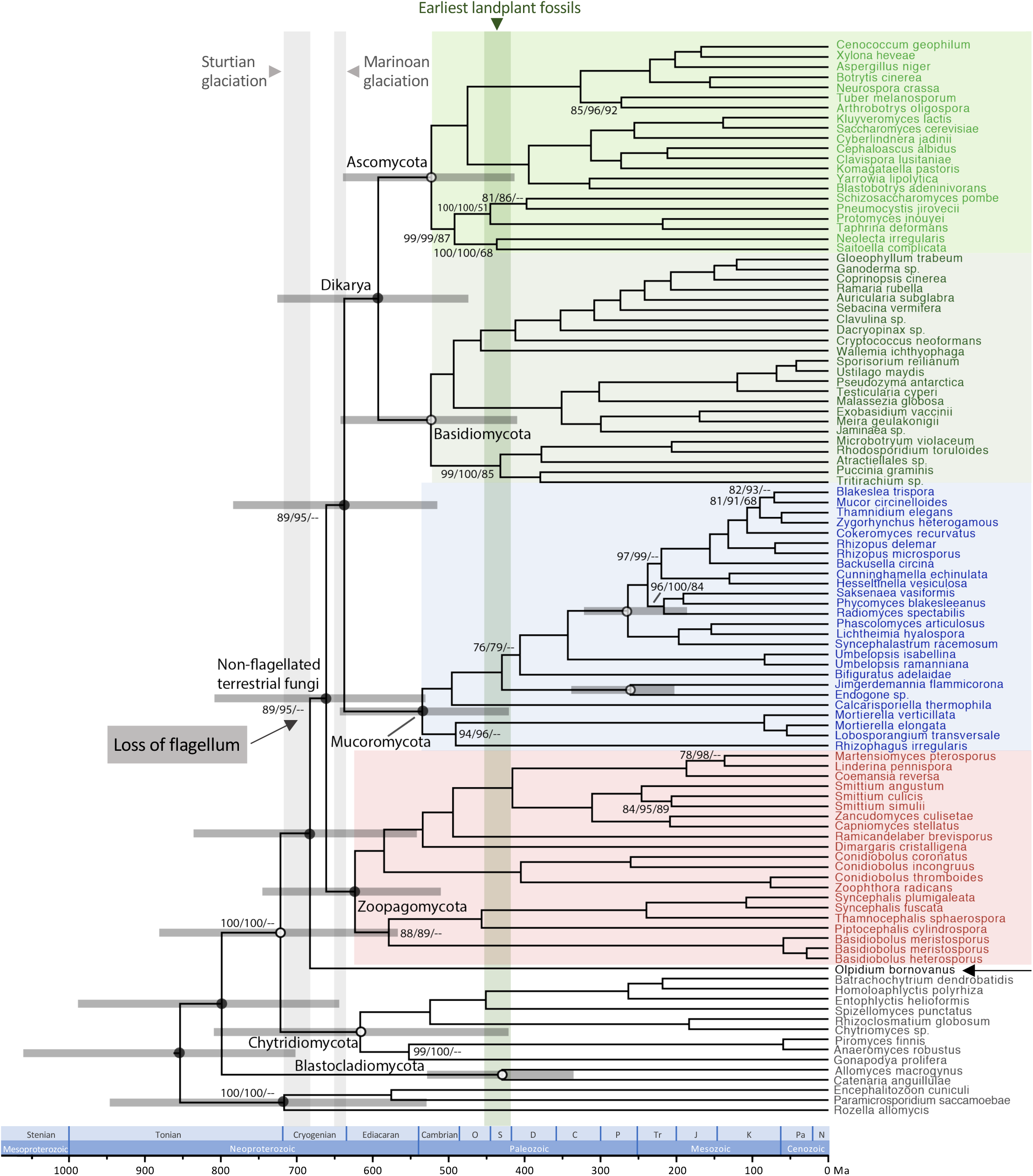
A dated phylogeny of Kingdom fungi. The chronogram is based on the best ML tree inferred with RAxML (113 taxa, 295 proteins and 83,461 aligned amino acid sites) and the divergence times are estimated with MCMCTree. The numbers by the nodes are the bootstrap support from three analyses (non-partitioned RAxML analysis, IQ-TREE analysis using heterotachy model and ASTRAL analysis) based on the 295-marker dataset. No bootstrap values are shown for nodes with support higher than 90 in all three analyses. Open circles denote the calibration points used in MCMCTree analysis (Table S4). The grey vertical shade indicates the durations of the Sturtian and Marinoan glaciations. The green vertical shade denotes the geological time that the earliest landplant fossils were from.

We further assessed the completeness of the dataset to evaluate whether missing data may impact the phylogenetic reconstruction. All the sampled markers were present in 80% of the sampled genomes, except for one that was found in 75% of all sampled genomes (85 genomes). All the genomes possessed at least 80% of all the 295 markers, except for the genome of *Encephalitozoon cuniculi* (Cryptomycota) with only 137 sampled markers present. Of all the 295 markers, 283 had broad phylogenetic distributions as defined by being present in *Olpidium* and at least one species of each of Zoopagomycota, Mucoromycota, Ascomycota and Basidiomycota. Phylogenetic reconstruction based on these 283 markers was

Four of the markers contained identical sequences from two or more different species in the individual alignments after trimming and were excluded in the RAxML inference of individual gene trees and the ASTRAL analyses ^34^. The topology recovered in the ASTRAL analysis based on all remaining 291 individual gene trees was mostly consistent with the ConcatBestTree (Fig. 2). One relevant discrepancy, however, was the placement of *Olpidium* and *Basidiobolus*. In the ASTRAL analysis based on the original individual gene tree, *Olpdium* and *Basidiobolus* formed a paraphyletic grade subtending Mucoromycota. When the weak nodes (bootstrap support <30) were collapsed for the individual gene trees, ASTRAL analysis recovered *Basidiobolus* as a member of Zoopagomycota and *Olpidium* as sister to Mucoromycota/Dikarya clade.

The dating analysis using MCMCTree ^35^ estimated the age of the MRCA of all fungi as 854 Ma (Fig. 2). The age of MRCA of *Olpidium* and the non-flagellated fungi was estimated as 683 Ma. The age of MRCA of all non-flagellated fungi was estimated as 661 Ma, while the split between Mucoromycota and Dikarya was estimated as 638 Ma. The ages of the MRCAs of each fungal phylum were estimated as 430 Ma for Blastocladiomycota, 617 Ma for Chytridiomycota, 624 Ma for Zoopagomycota, 535 Ma for Mucoromycota, 524 Ma for Basidiomycota and 523 Ma for Ascomycota, while the age of MRCA of Dikarya was estimated as 593 Ma (Fig. 2).

### Investigation of the uncertain local placement of *Olpidium* in fungal tree of life and the branching order between Blastocladiomycota and Chytridiomycota

RAxML analyses with fast-evolving sites removed from the concatenated 295-marker matrix recovered the same topology of (*Olpidium*, (Zoopagomycota, (Mucoromycota, Dikarya))), with a minor to moderate decrease of branch support compared with analyses included all sites (Table S3). We performed two sets of the Approximately Unbiased (AU)^36^ tests, one set on the concatenated matrices of 295 markers and the other set on individual markers (Fig. S1). For the AU tests on individual markers, a total of 283 markers were included from the analysis. Four of the 12 excluded markers contained identical sequences from two or more different species in the trimmed individual alignments. For the other eight excluded markers, one or more constraints used in RAxML analysis could not be applied to due to missing taxa and thus the AU test could not be performed on these markers.

The AU test on the concatenated 295-marker matrix was consistent with the ConcatBestTree being significantly better than the alternative topologies. For the AU tests on individual markers, the ConcatBestTree was not rejected as a significantly worse tree than the best individual gene trees for 236 markers (the ‘AU_similar’ set of markers), while the best gene trees for 47 markers were significantly better than the ConcatBestTree (the ‘AU_better’ set of markers) (Fig. S1). The remaining 12 markers were not eligible for AU test due to the presence of missing data. Out of the 236 markers in the AU_similar set, 188 of them showed equivalent support for at least one other alternative topology in addition to the best gene tree and the ConcatBest tree. (Fig. S1). In the quartet mapping analysis on the concatenated 295-marker matrix, 48.2% of the sampled quartet favored the topology of ((*Olpidium*, Mucoromycota), (Zoopagomycota, Dikarya)), 39.6% favored ((*Olpidium*, Zoopagomycota), (Mucoromycota, Dikarya)), and the remaining 11.4% favored ((*Olpidium*, Dikarya), (Mucoromycota, Zoopagomycota)). In the quartet mapping analysis on individual markers, only one marker had more than 75% of the sampled quartet favored the topology of ((*Olpidium*, Zoopagomycota), (Mucoromycota, Dikarya)), while all the remaining markers show no or little preferences over any of the three competing topologies (Fig. 3).

**Fig. 3.**
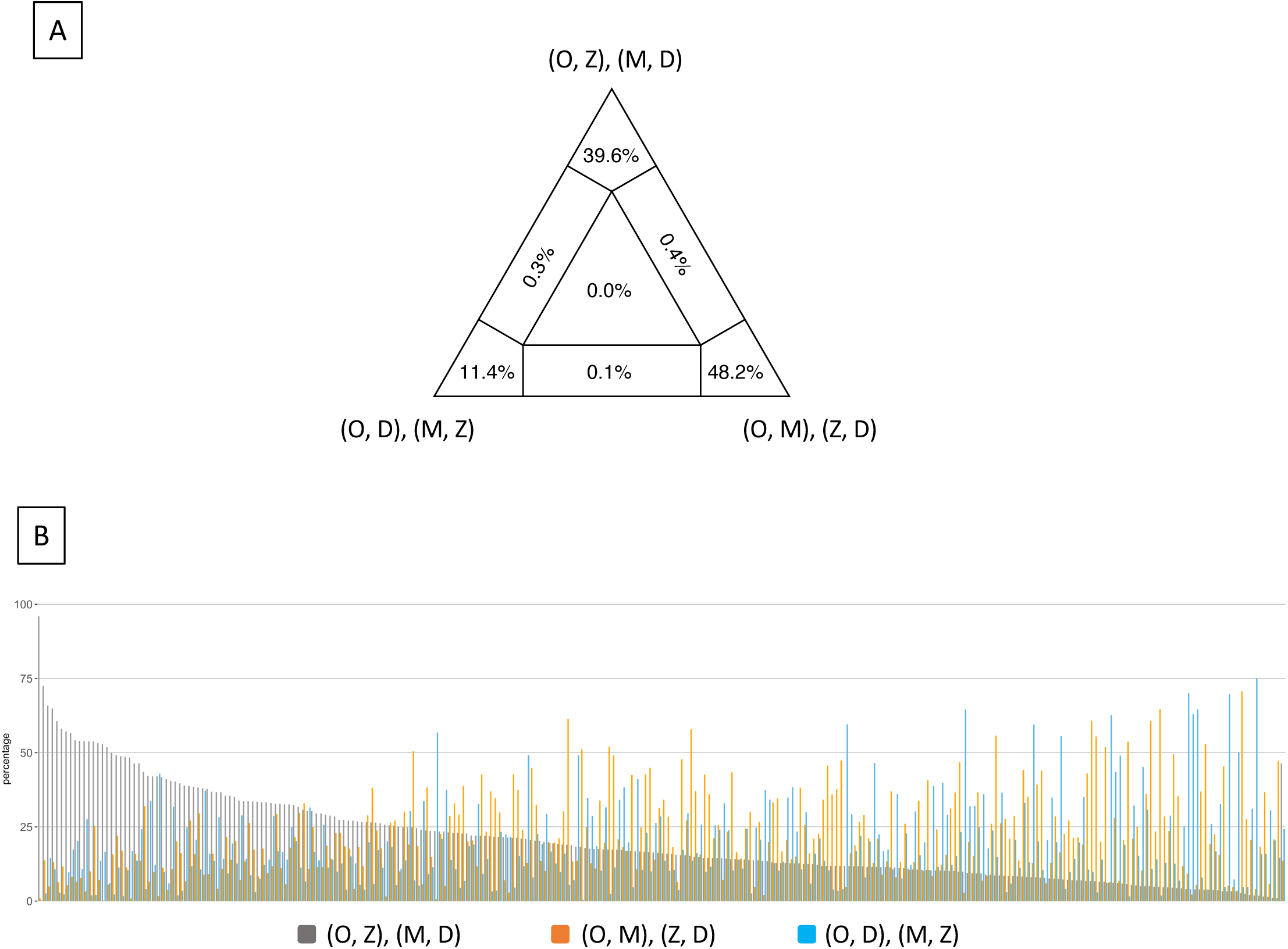
Quartet mapping analyses on the 295-marker dataset showed equivocal support for the competing hypotheses regarding the relationships among *Olpidium*, Mucoromycota, Zoopagomycota and Dikarya. A). Quartet mapping analysis based on the concatenated matrix represented as an equilateral triangle. Values at the corners indicate the percentages of quartets considered resolved over the total number of sampled quartets. Values at the lateral and central regions represent the percentages of unresolved quartets. B). Quartet mapping analyses on individual markers. Each marker is plotted with three bars with the height being the percentage of quartets considered resolved over the total number of quartets sampled for a particular marker. D = Dikarya, M = Mucoromycota, O = *Olpidium*, Z = Zoopagomycota.

To take the uncertainty of individual gene trees into account, we collapsed the weakly supported nodes (bootstrap support < 30) in the ASTRAL polytomy tests. In the polytomy test based on all the 291 individual gene trees, the length of the internode between the divergence of *Olpidium* and Zoopagomycota was reduced to zero (Fig. S1). The polytomy test based on the 236 markers from the AU_similar set identified a polytomy regarding the relationships among *Olpidium*, Zoopagomycota, Mucoromycota and Dikarya. For the 47 markers from AU_better set, there were insufficient quartets remaining to perform the polytomy test when the weak nodes in the gene trees (bootstrap support < 30) were collapsed. When the original gene trees were used, the polytomy among *Olpidium*, Zoopagomycota, Mucoromycota and Dikarya was again recovered.

We also performed AU tests on the individual gene trees on the relationships among Blastocladiomycota, Chytridiomycota and the clade of *Olpidium* and non-flagellated terrestrial fungi, which were also represented by a polytomy in tests by ASTRAL. We performed AU tests on the individual gene trees regarding the diverging order between Blastocladiomycota and Chytridiomycota. Among the 262 markers that were eligible for AU tests, 212 markers showed equivocal support for the individual best gene tree and for the branching pattern found in the best tree, i.e., (Blastocladiomycota, (Chytridiomycota, (*Olpidium*, non-flagellated terrestrial fungi))) (referred as Blasto_sister arrangement). The best gene trees of the remaining 50 markers differed from and were significantly more likely than the Blasto_sister arrangement.

## Discussion

### Generating DNA data for phylogenetic reconstruction for fungi present in complex biological systems

The obligate endoparasitic nature of *Olpidium* makes it challenging to collect DNA of the quantity and quality necessary for whole-genome sequencing. The fungus produced zoosporangia in the roots of inoculated cucumbers grown in sand. Transferring the roots with zoosporangia to distilled water triggered differentiation and release of zoospores into the water, but it also allowed dispersal of other organisms living within or on the surface of the host plant roots. Hence the genomic data generated from the *Olpidium* zoospores is a mixture of DNA sequences with different origins. We were able to retrieve *Olpidium* sequences with strong confidence by applying binning methods based on oligonucleotide compositions by VIZBIN and supplemented by BLAST and mmseq2 searches. While our conservative sampling approach in VIZBIN analysis might lead to the exclusion of some *Olpidium* sequences from the working assembly, it assures that the sequences included in the working assembly were highly likely from *Olpidium*. The quality of the working assembly was further boosted by the additional filtering of data with sequence-similarity-based methods of BLAST and mmseq2. Even though the size of the working assembly is only 16% of the raw assembly, the BUSCO score of the working assembly is 75%. Out of the 434-marker set that proved useful to reconstruct high-level fungal relationships ^37,38^, we were able to retrieve 295 of them for use in the subsequent phylogenetic analysis.

Generating adequate genomic data from non-culturable fungi represents a challenge in the incorporation of these organisms in phylogenetic analyses, especially for fungi that are microscopic for their entire life histories and/or reside in a complex biological system (e.g., endophytes, endoparasites, soil fungi etc.). Our approach provides a solution to effectively retrieve markers for phylogenomic inference from these problematic taxa without demanding extraordinary measures in acquisition of sequence data. It can also serve as an approach to survey the fungal diversity of complex systems (such as the microbiome of soil samples) and potentially perform functional analyses of these systems, especially with the help of the rapid advancement of binning methods targeting eukaryotic sequences ^28^.

### *Olpidium* is the sister group to non-flagellated, terrestrial fungi

Loss of the flagellated stage is considered to be one of the most important morphological transitions in the evolution of fungi and has been interpreted to be associated with the colonization of the terrestrial landscape and rise of the non-flagellated fungi ^39,40^. Being a zoospore-producing fungus, *Olpidium* was long considered a member of Chytridiomycota, but multi-gene studies by James *et al*.^2^ and Sekimoto *et al*.^16^ questioned this hypothesis. These analyses grouped *Olpidium* within the non-flagellated terrestrial fungi, but taxon sampling was relatively sparse and the exact phylogenetic placement of *Olpidium* remained unresolved. In this study we densely sampled within ‘zygomycete’ fungi, i.e., Mucoromycota and Zoopagomycota ^2^, aiming to include the closest extant relatives of *Olpidium*. We also sampled multiple species from each of the individual classes within Dikarya and within Chytridiomycota to capture the deepest nodes, i.e., the MRCAs of all extant phyla and subphyla, within these two groups, allowing us to break potential long branches between these major fungal lineages.

Our analyses based on the super-alignment approach (RAxML and IQ-TREE) inferred *Olpidium* as more closely related to the non-flagellated terrestrial fungi, consistent with the findings by James *et al*.^15^ and Sekimoto *et al*. ^16^. In contrast to these two studies where *Olpidium* was found nested within non-flagellated fungi, however, *Olpidium* was recovered as the sister group to the clade of non-flagellated terrestrial fungi. This topology supports the most parsimonious model of morphological evolution with the production of zoospores through zoosporangia being a symplesiomorphic character shared by *Olpidium* and the ‘core chytrids’, and a single loss of the flagellum leading to the MRCA of (Zoopagomycota, (Mucoromycota, Dikyara)), rather than multiple losses among the fungi now classified in Zoopagomycota.

### Lack of strong support for recalcitrant nodes along the fungal backbone is due to lack of phylogenetic information

The analyses performed here do not support missing data as being a major contributing factor to the uncertain placement of *Olpidium* relative to the non-flagellated terrestrial fungi. The removal of fast-evolving sites consistently supported the branching order of (*Olpidium*, (Zoopagomycota, (Mucoromycota, Dikarya))) with minor to moderate decrease of branch support (Table S3), suggesting that the noise potentially brought in by fast-evolving sites was limited in the inference of the placement of *Olpidium*. Quartet analysis, AU tests, and direct examination of individual gene trees revealed little conflict among the sampled markers. Only eight markers showed strong support for an alternative placement of *Olpidium*. The removal of these eight markers from the 295-marker dataset did not improve the support for the placement of *Olpidium* in the subsequent phylogenetic reconstruction, suggesting that these conflicting signals were not the major contributing factor to the uncertain local placement of *Olpidium* (Table 1). When individual markers were ranked based on their phylogenetic informativeness (PI) and added to a concatenated dataset in a stepwise fashion from highest to lowest PI, the node supporting *Olpidium* with the non-flagellated fungi was strongly supported and stably resolved with twenty markers (Table 1). Resolution of the relationships between Zoopagomycota, Mucoromycota and Dikarya required more than 200 genes, however, and never reached maximum values of support, a finding consistent with limited phylogenetic signal.

**Table 1.** Summary of branch support regarding the placement of *Olpidium* and the main non-flagellated terrestrial fungal groups in RAxML analyses with the markers added sequentially according to their informativeness. For each dataset, we performed RAxML analysis with 100 bootstrap replicates and PROTGAMMALG model. The arrangement of (O, (Z, (M, D))) was recovered in all the RAxML analyses. Bootstrap higher than 50 and posterior probabilities higher than 0.5 are shown. D – Dikarya; M – Mucoromycota; O – *Olpidium bornavanus;* Z – Zoopagomycota.

| # of markers included | 5 | 10 | 20 | 30 | 40 | 50 | 60 | 70 | 80 | 90 | 110 | 130 | 150 | 170 | 190 | 210 | 230 | 250 | 270 | 290 |
| --- | --- | --- | --- | --- | --- | --- | --- | --- | --- | --- | --- | --- | --- | --- | --- | --- | --- | --- | --- | --- |
| (O, Z, M, D) | -- | -- | -- | 81 | 92 | 100 | 100 | 100 | 100 | 100 | 100 | 100 | 100 | 100 | 100 | 100 | 100 | 100 | 100 | 100 |
| (Z, M, D) | -- | -- | 99 | -- | 63 | 85 | 96 | 87 | 76 | 52 | 63 | 61 | 67 | 48 | 73 | 67 | 77 | 92 | 93 | 96 |
| (M, D) | -- | -- | -- | -- | -- | 84 | 96 | 86 | 76 | 52 | 63 | 61 | 66 | 48 | 72 | 67 | 77 | 93 | 93 | 96 |

| # of markers included | 5 | 10 | 20 | 30 | 40 | 50 | 60 | 70 | 80 | 90 | 110 | 130 | 150 | 170 | 190 | 210 | 230 | 250 | 270 | 287 |
| --- | --- | --- | --- | --- | --- | --- | --- | --- | --- | --- | --- | --- | --- | --- | --- | --- | --- | --- | --- | --- |
| (O, Z, M, D) | -- | -- | 100 | 100 | 100 | 100 | 100 | 100 | 100 | 100 | 100 | 100 | 100 | 100 | 100 | 100 | 100 | 100 | 100 | 100 |
| (Z, M, D) | -- | -- | -- | -- | -- | 65 | 51 | -- | -- | -- | -- | -- | -- | 61 | -- | 55 | 62 | 76 | 82 | 79 |
| (M, D) | -- | -- | -- | -- | -- | 64 | 50 | -- | -- | -- | -- | -- | -- | 61 | -- | 55 | 62 | 76 | 82 | 79 |

The equivocal support in AU tests and quartet analyses for three or more competing phylogenetic positions of *Olpidium* suggests that limited phylogenetic information contained in the markers is the main cause of the lack of strong support for the nodes defining the relationships between *Olpidium*, Zoopagomycota and Mucoromycota. Similarly, AU tests on the placement of Blastocladiomycota and Chytridiomycota also suggested lack of phylogenetic signals in individual markers. Consistent with the AU tests, the ASTRAL polytomy tests on the super-tree datasets recovered a polytomy for the branching order among *Olpidium*, Zoopagomycota, Mucoromycota, and Dikarya. These findings are consistent with the branches involved in these ASTRAL polytomies being one-third to one-half the lengths of the flanking backbone branches (Fig. 2). These short branches would be best captured by more quickly evolving genes/sites, however, the ancient age of these evolutionary events further compounds this problem and creates a situation that the genes/sites with the most appropriate substitution rate are also more susceptible to signal decay over time.

### The loss of the flagellum in fungi and the radiation of the early terrestrial fungi

The MRCAs of the Chytridiomycota, Zoopagomycota, Mucoromycota and Dikarya are all estimated to be of similar and overlapping ages (Fig. 2), lending support to the hypothesis of rapid radiation among the early terrestrial fungal groups ^16,41^. These findings are consistent with a Precambrian origin of the non-flagellated fungi ^5,42,43^, and that they experienced their first radiation in Neoproterozoic (Fig. 2), concurrent with Ediacaran fauna (625-539 Ma)^44^ and predating the earliest evidence of land plants in the late Ordovician and Silurian (450-420 Mya)^45^. While animal phyla likely diversified in early marine environments, the loss of flagellum and concurrent ages of the MRCA of Zoopagomycota, Mucoromycota and Dikarya suggest that these lineages likely diversified in early terrestrial environments.

The Neoproterozoic was an era characterized by multiple glaciation events, commonly referred to as the ‘snowball’ earth^46^. Under this model, the MRCA of the non-flagellated terrestrial lineages and its ancestors must have resided in the microbial biocrust, which has been proposed as the earliest terrestrial ecosystem ^47–49^. These early terrestrial fungi would have possessed a zoosporic stage after they first moved to the microbial biocrust, and the glaciations, especially the Sturtian glaciation, might have led to the extinction of much of the diversity. The survivors of the glaciation rapidly radiated during the inter-glaciation period or post-glaciation early Ediacaran, which was characterized by increased temperatures, newly created terrestrial niches created by the retreating glaciers, and new food sources by the diversification of other eukaryotes (e.g., protists and algae) ^50–52^.

The loss of the flagellum might have either occurred during the glaciations, or before fungi began to diversify on land after the glaciation. In either scenario, *Olpidium* and other zoosporic species retained their flagella and remained dependent on liquid for dispersal. Whereas the remaining terrestrial fungal lineage lost the flagellated zoospore stage and invented means to disperse spores aerially, which freed them from dependence on free water for dispersal. In addition, hyphal growth form of these fungi allowed them to efficiently explore the terrestrial landscape. As a result, these non-flagellated fungi radiated and gave rise to 1.5 million species and became the dominant fungi in terrestrial, plant-based ecosystems.

## Methods

### Genome and transcriptome sequencing and assembly

*Olpidium bornovanus* S191 was co-cultured with cucumber roots in sand. At maturity, the zoospores of *O. bornovanus* were collected by washing the cucumber root with sterile water. The collected zoospores were pelleted at 5,000 × g for five minutes. Total DNA and RNA were extracted from the pelleted zoospores using Qiagen DNeasy Plant Mini kit and Qiagen RNeasy Plant Mini kit, respectively, following the procedures outlined by the manufacturer.

The *Olpidium* genome was sequenced using both mate-pair sequencing and standard Illumina pair-end sequencing. For mate-pair sequencing, the amplified paired-end library was size selected to obtain the desired fragments (300 bp – 600 bp) and sequenced on the Illumina HiSeq2000 sequencer. The paired-end library was sequenced on an Illumina HiSeq2000, following a 2×100 indexed run recipes. The *Olpidium-*raw-genome assembly was generated using MEGAHIT ^53^ based on raw reads from both the paired end and mate pair sequencing runs.

To prepare the library for transcriptome sequencing, messenger RNA was purified from total RNA using Dyanbeade mRNA isolation kit (Invitrogen) and was fragmented using RNA Fragmentation Reagents (Ambion) with targeting fragments range of 300bp. Complementary DNA was generated by reverse transcription of the fragmented DNA using SuperScript II Reverse Transcription (Invitrogen). The cDNA library was prepared was sequenced using Illumina HiSeq2000 sequencing, following a 2×150 indexed run recipe. The raw RNA-Seq reads were assembled using TRINITY v.2.5.1^54^.

### Binning analysis on *Olpidium* transcriptome and genome assemblies

*Olpidium* transcriptome contigs longer than 1.5 kb were binned based on tetramer composition of individual contigs using VIZBIN. We identified a total of seven clusters of contigs (Fig. 1A). Contigs in each data cluster were searched against Refseq database ^55^ and an in-house database composed of 350 fungal proteomes. Cluster 1 (Fig. 1A) was dominated by sequences showing fungal affinities and was used in the subsequent binning analysis on the raw genome assembly of *Olpidium* (*Olpidium-*raw-genome).

In the second VIZBIN analysis, we included contigs from the *Olpidium-*raw-genome assembly and contigs from cluster 1 of the *Olpidium* transcriptome assembly. We subsampled the contig cluster of *Olpidium-*raw-genome contigs that overlapped with the majority of transcriptome cluster 1 contigs (dataset *Olpidium*-sub1; Fig. 1B).

*Olpidium*-raw-genome contigs longer than 1kb were searched against the in-house database of fungal proteomes using DIAMOND blastx ^56^. The genome sequences of the two fungi with the highest number of hits among Dikarya taxa (*Coprinopsis cinereal* and *Ustilago hordei*), together with genome sequences from two Mucoromycota fungi (*Mortierella elongata* and *Umbelopsis ramanniana*,), two Zoopagomycota fungi (*Basidiobolus meristosporus* and *Syncephalis plumigaleata*), and one Chytridiomycota fungus (*Spizellomyces punctatus*), together with contigs from *Olpidium*-sub1, were included in a third VIZBIN binning analysis. Prior to the third binning analysis, the aforementioned seven fungal genome sequences were divided into 3-kb fragments using a custom script. We removed the contigs from *Olpidium*-subsample1 that clustered with the non-*Olpidium* reference genome sequences. The remaining *Olpidium*-subsample1 (dataset *Olpidium*-subsample1’; Fig. 1C) was used to query the NCBI nr and Mycocosm database using mmseq2 ^57^. Each contig with the top hit from a non-fungal organism was extracted and compared against an in-house database composed of 350 fungal proteomes using DIAMOND with the blastx option. The contigs without significant fungal hits (1e-30) in the DIAMOND search were removed from *Olpidium*-subsample1’. The remaining contigs from *Olpidium*-subsample1’ were used as the working assembly of *Olpidium*.

### Genome annotation

Prediction of protein-coding genes in the working assembly of *Olpidium* was performed using MAKER pipeline v2.31.8^58^. The genome was masked with RepeatMasker v 4-0-7) (Smit, AFA, Hubley, R & Green, P. RepeatMasker Open-4.0. 2013-2015 http://www.repeatmasker.org) with the species set to “fungi” from RepBase^59^. Gene prediction was performed with the ab initio gene predictors SNAP v2017-03-01^60^, GeneMark.HMM-ES v4.33^61^, and Augustus v3.3 (rhizopus_oryzae model)^62^. Prediction parameters for Augustus and SNAP both used Rhizopus oryzae trained models. Transcript evidence for genome annotation was generated from the RNA-Seq reads assembled using Trinity v.2.5.1^54^. Prediction of tRNA genes was performed with tRNAScan-SE v1.3.1^63^.

### Phylogenetic reconstruction and molecular dating analysis

We sampled proteome sequences from a total of 113 species (106 fungi and seven non-fungus outgroups; Table S1) in our phylogenetic reconstruction. We employed 434 protein markers that proved useful for the inference of higher-level phylogenetic relationships among fungi ^29,38^ (https://github.com/1KFG/Phylogenomics_HMMs). We used the pipeline PHYling (Stajich JE; http://github.com/stajichlab/PHYling_unified) to search for the target markers in the sampled proteomes with an *e-*value of −20 and performed alignment for each marker. A total of 295 targeted markers were found present in the working assembly of *Olpidium* and these markers were the main focus in the subsequent analyses.

We took three approaches to infer the phylogenetic relationships of the sampled taxa: the super-alignment approaches by using RAxML v.8.0.26 and IQ-TREE on the concatenated alignment, and the super-tree approach by using ASTRAL-III on the individual markers. The RAxML analysis on the concatenated matrix was performed on the CIPRES web portal ^64^, using the “-f a” option with 100 bootstrap replicates. We performed non-partitioned analysis with PROTGAMMALG model and partitioned analysis using a partition scheme generated by PartitionFinder2 ^65^. For IQ-TREE analyses, the best ML tree was based on the non-partitioned 295-marker dataset, using a FreeRate heterogeneity model (LG+F+R10) and using a heterotachy model (LG+F+R10*H4) ^66,67^. Branch supports for IQ-TREE analyses were estimated using the ultrafast bootstrap approximation approach with 1000 bootstrap replicates (Hoang *et al*., 2018). For the ASTRAL analysis, we initially scored the topology obtained in the RAxML analysis of the concatenated alignment (ConcatBestTree) with individual gene trees. We then inferred the best ASTRAL topology based on individual gene trees. To account for the uncertainty in the best gene tree, we collapsed the weakly supported nodes (bootstrap support < 30) in the individual gene trees. We also performed a bootstrap analysis (100 replicates) and generated a greedy consensus of the 100 bootstrap replicates.

Molecular dating analyses were performed on the best RAxML tree based on the 295-marker matrix using MCMCTree of the PAML4.9 package, with only the fungal ingroups included. We used six fossils as the lower minimum bound for the MRCAs of Blastocladiomycota (407 Ma), Chytridiomycota (407 Ma), Endogonaceae (247 Ma), Mucorales (315 Ma), Ascomycota (407 Ma) and Basidiomycota (330 Ma) (Table S4). A truncated Cauchy distribution was applied for each of the calibration points. In addition, we set a loose maximum age constraint on the node of the MRCA of Chytridiomycota, *Olpidium* and the non-flagellate terrestrial fungi (MRCA-COT) based on expansion of fungal pectinases and the age of pectin-containing streptophytes as discussed in Chang *et al*.^5^. The recent estimates on the origin of streptophytes range from 750 Ma to about 1100 Ma and the origin of the pectin-containing lineages were around 850 Ma or younger^43,69,70^. We applied 850 Ma as the upper bound of MRCA-COT. We used a relaxed molecular clock with the independent rate model (clock = 2) ^71^, and divergence times were estimated using the approximation method ^72^. We performed two independent MCMC runs with the following parameters: burn in = 250,000; sampling frequency = 100; number of samples = 20,000. With this setting, the first 250,000 iterations were thus discarded as burn-in, and then the MCMC was run for another 2,000,000 iterations, sampling every 100 iterations. The 20,000 samples were then summarized to estimate mean divergence date and associated 95% credibility intervals.

### Investigation of the uncertainty in the local placement of *Olpidium* in fungal tree of life

To explore the impact of fast-evolving sites on the phylogenetic reconstruction, we used TIGER 1.02 ^73^ to classify all the aligned amino acid sites into ten rate categories. We sequentially removed the sites with faster substitution rates from the matrix and performed tree inference using RAxML with 100 bootstrap replicates.

To investigate whether the lack of strong support for the local placement *Olpidium* in Kingdom Fungi was due to conflicting signals from individual genes or due to lack of informative sites, we performed the Approximately Unbiased (AU) tests and the polytomy tests implemented in ASTRAL(Fig. 2). We performed AU tests on the concatenated matrix based on the 295-marker dataset and on the individual markers. We identified the alternative placements of *Olpidium* present in the individual RAxML bootstrap trees based on the 295-dataset and in the ASTRAL analysis. We applied each of the *Olpidium* placements as topology constraints in RAxML analyses to generate the alternative trees used in AU tests. For AU tests on individual markers, we applied an additional ConcatBestTree constraint with *Olpidium* being sister to the non-flagellated terrestrial fungi. For each AU test, the site likelihood was calculated using RAxML and the final AU test was performed using CONSEL^74^. The ASTRAL polytomy test was applied to the concatenated matrices and to subsets of the matrices which were generated based on the results of AU tests (Fig. S1). For each dataset we performed one test on the original gene trees and one test on the modified gene trees with their low-supported nodes collapsed (bootstrap value < 30) to account for the uncertainty of individual gene trees.

To further explore the support for the relationships among *Olpidium*, Zoopagomycota, Mucoromycota and Dikarya, quartet mapping analyses were performed in IQ-TREE using the options -lmclust to define the aforementioned four groups. Additionally, phylogenetic informativeness of the 295 loci was measured using PhyDesign web server ^75,76^ (http://phydesign.townsend.yale.edu/). Phylogenetic informativeness (PI) was estimated from the concatenated and partitioned 295-marker dataset and the chronogram generated from MCMCTREE. We ranked the 295 markers according to their phylogenetic informativeness and sequentially added the marker to a series of concatenated matrices according to their PI rank from highest to lowest. We inferred phylogenetic trees based on each of the resulting matrices using IQ-TREE.

## Data Availability Statement

The genome and transcriptome data generated in this study is in the process of depositing to GenBank. The accession numbers will soon be available and included in the revised manuscript.

## Acknowledgements

This material is based upon work supported by the National Science Foundation grant DEB- 1441604 to JWS. JES is a CIFAR Fellow in the program Fungal Kingdom: Threats and Opportunities. YW and JES were supported by NSF grants DEB-1441715 and DEB-1557110. The work conducted by the U.S. Department of Energy Joint Genome Institute, a DOE Office of Science User Facility, is supported by the Office of Science of the U.S. Department of Energy under Contract No. DE-AC02-05CH11231. Any opinions, findings, and conclusions or recommendations expressed in this material are those of the author(s) and do not necessarily reflect the views of the National Science Foundation. The authors are grateful to Drs. G. Bonito, C. Cuomo, T. James, and R. Vilgalys for giving the permission to use their unpublished genome data. We would also like to thank Dr. Stephen Mondo for his help on data preparation for the submission to JGI MycoCosm and NCBI. Special thanks are given to Dr. Mary Berbee for her coordination on the initial *Olpidium* genome project and the constructive discussion and comments during the preparation of the manuscript.

## Author contributions

Y.C. and J.W.S. conceived and designed the data analysis. D.R. and S.S. performed DNA & RNA material collection. M.C. and L.S. carried out genome and transcriptome sequencing. J.E.S. performed genome and transcriptome assembly and annotation. A.S contributed to metagenome binning. Y.C. performed data analysis on metagenome binning and phylogenetic investigation. Y.C. and J.W.S wrote the draft of manuscript. Y.W., I.V.G. and J.E.S. contributed to manuscript revision.

## Figure Legends

Figure S1. The flowchart showing the procedure and results of phylogenetic reconstruction, topology tests and polytomy tests.

