## Supplementary for "Genome-scale phylogenetic analyses confirm *Olpidium* as the closest living zoosporic fungus to the non-flagellated, terrestrial fungi"

Fig. S1. The flowchart showing the procedure and results of phylogenetic reconstruction, topology tests and polytomy tests.

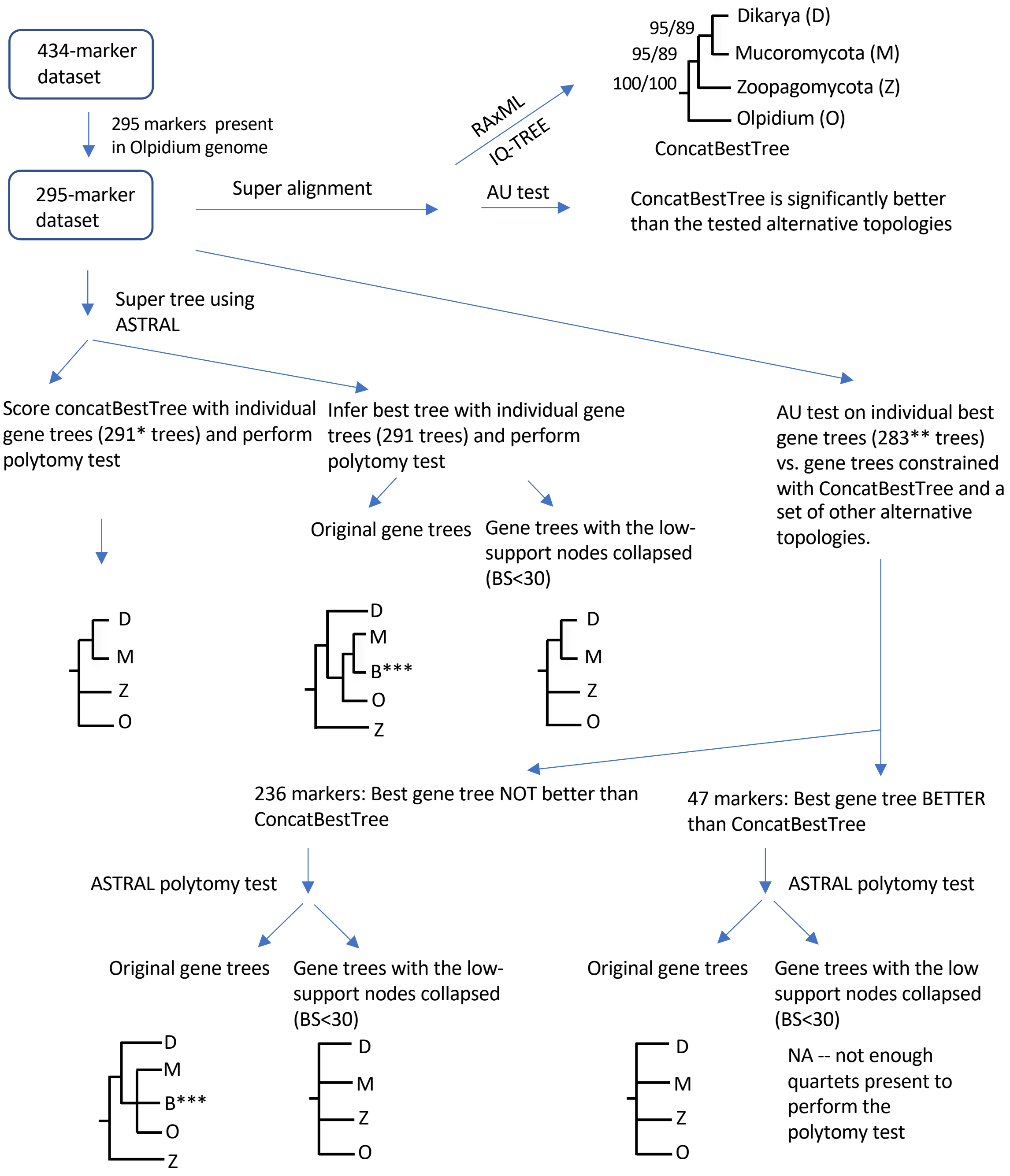


*: Four individual marker alignments contain identical sequences from two different species or contain sequence(s) with only undetermined values; hence no RAxML analysis on those markers.
**: No constrained-tree search is done for four markers due to the large number of missing species for those markers. Together with the four markers with all-undetermined sequences/identical sequences for two different species, a total of eight markers are not eligible for AU test.
***: B stands for *Basidiobolus*.

Table S1. Genome information of the 113 taxa included in the phylogenetic reconstruction.

| Species | GenBank accession No./JGI Web Portal (reference) |
|---|---|
| *Allomyces macrogynus* ATCC 38327 | ACDU00000000.1 |
| *Anaeromyces robustus* | https://genome.jgi.doe.gov/Anasp1/ [1] |
| *Arabidospsis thalina* | GCA_000001735.2 [2] |
| *Arthrobotrys oligospora* ATCC 24927 | ADOT00000000 [3] |
| *Aspergilus fumigaus* ATCC 1015 | ACJE00000000 [4] |
| *Atractiellales* sp. | https://mycocosm.jgi.doe.gov/Atrsp2/ |
| *Auricularia subglabra* | https://mycocosm.jgi.doe.gov/Aurde3_1/A [5] |
| *Backusella circina* FSU 941 | http://genome.jgi.doe.gov/Bacci1 |
| *Basidiobolus heterosporus* B8920 | GCA_000697455.1 [6] |
| *Basidiobolus meristosporis* B9252 | GCA_000697375.1 [7] |
| *Basidiobolus meristosporis* CBS 931.73 | https://mycocosm.jgi.doe.gov/Basme2finSC/ [7] |
| *Batrachochytrium dendrobatidis* JAM81 | ADAR00000000.1 |
| *Bifiguratus adelaidae* AZ0501 | MVBO00000000 [8] |
| *Blakeslea trispora* NRRL 2456 | https://mycocosm.jgi.doe.gov/Blatri1/ |
| *Blastobotrys adeninivorans* LS3 | https://mycocosm.jgi.doe.gov/Arxad1/ [9] |
| *Botrytis cinerea* B05.10 | GCA_00143535.4 [10] |
| *Calcarisporiella thermophila* CBS 279.70 | https://gb.fungalgenomics.ca/portal/ |
| *Capniomyces stellatus* MIS-10-108 | LUVW010000000 [11] |
| *Capsaspora owczarzaki* ATCC 30864 | ACFS00000000.2 [12] |
| *Catenaria anguillulae* PL171 | http://genome.jgi.doe.gov/Catan1 [7] |
| *Cenococcum geophilum* 1.58 | https://mycocosm.jgi.doe.gov/Cenge3/ [13] |
| *Cephaloascus albidus* ATCC 66658 | https://mycocosm.jgi.doe.gov/Cepfr1_1 |
| *Chlamydomonas reinhardtii* CC-503 | ABCN00000000.2 [14] |
| *Chytriomyces* sp. MP 71 | https://mycocosm.jgi.doe.gov/Chytri1/ |
| *Clavispora lusitaniae* ATCC 42720 | AAFT00000000.1 [15] |
| *Clavulina* sp. PMI 390 | https://mycocosm.jgi.doe.gov/ClaPMI390 |
| *Coemansia reversa* NRRL 1564 | JZJC00000000 [16] |
| *Cokeromyces recurvatus* B5483 | https://mycocosm.jgi.doe.gov/Cokrec1/ |
| *Conidiobolus coronatus* NRRL 28638 | JXYT00000000 [16] |
| *Conidiobolus incongruous* B7586 | GCA_000697335.1 [16] |
| *Conidiobolus thromboides* FSU 785 | http://genome.jgi.doe.gov/Conth1 |
| *Coprinopsis cinerea* Okayama7_130 | AACS00000000.2 [17] |
| *Cryptococcus neoformans* JEC21 | GCA_000149245.3 [18] |
| *Cunninghamella echinulata* NRRL 1832 | https://mycocosm.jgi.doe.gov/Cunech1/ |
| *Cyberlindnera jadinii* NRRL Y-1542 | https://mycocosm.jgi.doe.gov/Cybja1/ [19] |
| *Dacryopinax* sp. DJM-731 | AEUS00000000.1 [5] |
| *Dictyostelium discoideum* | AAFI00000000.2 [20] |
| *Dimargaris cristalligena* RSA 468 | https://mycocosm.jgi.doe.gov/DimcrSC1/ [21] |
| *Drosophila melanogaster* vr6.04 | http://flybase.org [22] |
| *Encephalitozoon intestinalis* ATCC 50506 | GCA_000146465.1 [23] |
| *Endogone* sp. FLAS F-59071 | RBNK00000000.1 [24] |

| Species | GenBank accession No./JGI Web Portal (reference) |
|---|---|
| *Entophlyctis helioformis* JEL 805 | https://mycocosm.jgi.doe.gov/Enthel1/ |
| *Exobasidium vaccinia* MPITM | PRJNA196015 |
| *Funneliformis mosseae* DAOM-236685* | https://github.com/zygolife/AMF_Phylogenomics [25] |
| *Ganoderma* sp. 10597 SS1 | https://mycocosm.jgi.doe.gov/Gansp1/ [26] |
| *Gloeophyllum trabeum* ATCC 11539 | GCA_000344685.1 [5] |
| *Gonapodya prolifera* JEL478 | LSZK00000000 (Chang et al. 2015) [16] |
| *Hesseltinella vesiculosa* NRRL 3301 | http://genome.jgi.doe.gov/Hesve2finisherSC [7] |
| *Homoloaphlyctis polyrhiza* JEL142 | AFSM01000000.1 [27] |
| *Jimgerdemmannia flammicorona* AD002 | https://mycocosm.jgi.doe.gov/Jimfl_AD_1/ [24] |
| *Jimgerdemmannia flammicorona* GMNB39 | https://mycocosm.jgi.doe.gov/Jimfl_GMNB39_1/ [24] |
| *Jimgerdemmannia lactiflua* OSC 162217 | https://mycocosm.jgi.doe.gov/Jimlac1/ [24] |
| *Lichtheimia corymbifera* FSU 9682 | CBTN000000000.1 [28] |
| *Lichtheimia hyalospora* FSU 10163 | http://genome.jgi.doe.gov/Lichy1 |
| *Linderina pennispora* ATCC 12442 | http://genome.jgi.doe.gov/Linpe1 [7] |
| *Martensiomyces pterosporus* CBS 209.56 | http://genome.jgi.doe.gov/Marpt1 |
| *Monosiga brevicolis* MX1 | ABFJ00000000.1 [29] |
| *Mortierella elongata* AG-77 | http://genome.jgi.doe.gov/Morel2 [30] |
| *Mortierella verticillata* NRRL 6337 | AEVJ00000000.1 |
| *Mucor circinelloides* CBS 277.49 | http://genome.jgi.doe.gov/Mucci2 [31] |
| *Neurospora crassa* OR74A | AABX00000000.3 [32] |
| *Orpinomyces sp.* C1A | ASRE00000000.1 [33] |
| *Pandora formicae** | GCRV00000000.1 [34] |
| *Paraglomus brasilianum* DAOM-240472* | https://github.com/zygolife/AMF_Phylogenomics [25] |
| *Phycomyces blakesleeanus* NRRL 1555 | http://genome.jgi.doe.gov/Phybl2 [35] |
| *Piptocephalis cylindrospora* RSA 2659 | http://genome.jgi.doe.gov/Pipcy2/Pipcy2.home.html [21] |
| *Piromyces* sp. E2 | http://genome.jgi.doe.gov/PirE2_1 [1] |
| *Puccinia graminis f. sp. tritici* CRL 75-36-700-3 | AAWC0000000.1 [36] |
| *Racocetra castanea* BEG-1* | https://github.com/zygolife/AMF_Phylogenomics [25] |
| *Ramicandelaber brevisporus* CBS 109374 | http://genome.jgi.doe.gov/Rambr1 |
| *Rhizophagus diaphanous* MUCL 43196 | http://genome.jgi.doe.gov/Rhidi1 (Morrin *et al.*, 2019) |
| *Rhizophagus irregularis* DAOM 181602 | JARB00000000.1 [38] |
| *Rhizopus delemar* RA99-880 | AACW0000000.2 [39] |
| *Rhizopus microsporus var chinesis* CCTCCM201021 | CCYT00000000.1 [40] |
| *Rhizopus microsporus var microsporus* ATCC 52813 | http://genome.jgi.doe.gov/Rhimi1_1 [7] |
| *Rozella allomycis* CSF55 | ATJD00000000.1 [41] |
| *Saccharomyces serevisiae* S288C.vR642-1 | http://yeastgenome.org/ [42] |
| *Saksenaea vasiformis* B4078 | JNDT00000000.1 [6] |
| *Schizosaccharomyces pombe* 972h-.vASM294 | http://www.pombase.org/ [43] |
| *Scutellospora calospora* INVAM-IL209* | https://github.com/zygolife/AMF_Phylogenomics [25] |
| *Spizellomyces punctatus* DAOM BR117 | ACOE00000000.1 [44] |
| *Umbelopsis ramanniana* NRRL 5844 | http://genome.jgi.doe.gov/Umbra1 |
| *Ustilago maydis* 521 v190413 | AACP00000000.2 [45] |

| Species | GenBank accession No./JGI Web Portal (reference) |
|---|---|
| *Yarrowia lipolytica* CLIB 122 | GCA_000002525.1 [46] |
| *Zoophthora radicans* ATCC 208865 | http://genome.jgi.doe.gov/ZooradStandDraft_FD/ |

* Taxa with only transcriptome data.

Table S2. Summary of branch support regarding the placement of *Olpidium* and the main non-flagellated terrestrial fungal groups. All the ML-based analyses (i.e., RAxML and IQ-TREE analyses) recovered the same branching patterns in their best tree, (O, (Z, (M, D))), while (((O, (B, M)), D), Z') and ((B,(M, D)), Z') were the optimal topologies inferred by the ASTRAL analyses based on original gene trees and on gene trees with weak branches collapsed (BS < 30), respectively. Bootstrap higher than 50 and posterior probabilities higher than 0.5 are shown. B – *Basidiobolus*; D – Dikarya; M – Mucoromycota; O – *Olpidium bornavanus;* Z – Zoopagomycota; Z' -- Zoopagomycota without *Basidiobolus.*

| Analysis | RAxML | | IQ-TREE | | ASTRAL | |
|---|---|---|---|---|---|---|
| --no. markers | 295 | 295 | 295 | 295 | 295 | [1]295 |
| --models | LG+G | LG+G | LG+F+R10 | LG+F+R10*H4 | [2]LG+G | [2]LG+G |
| --other settings | partitioned | Non-partitioned | non-partitioned | non-partitioned | [3]original trees | [4]BS<30 collapsed |
| (O, Z, M, D) | 100 | 100 | 100 | 100 | 100 | 1 |
| (Z, M, D) | 66 | 89 | 96 | 95 | -- | -- |
| (M, D) | 68 | 89 | 96 | 95 | -- | -- |
| (O, M, B) | -- | -- | -- | -- | 82 | -- |
| (M, B) | -- | -- | -- | -- | 83 | -- |
| (O, M, D) | -- | -- | -- | -- | -- | 0.85 |

[1]Branch support shown is local posterior probabilities, computed based on a transformation of the percentage of quartets in individual gene trees that agree or disagrees with a branch.
[2]LG+G was the substitution model used in the inference of individual gene trees using RAxML.
[3]The original best individual gene trees were used in the ASTRAL analysis.
[4]In this ASTRAL analysis, the weakly supported branches (bootstrap < 30) were collapsed for each best individual gene tree.

Table S3. Summary of branch support regarding the placement of *Olpidium* and the main non-flagellated terrestrial fungal groups in RAxML analyses with faster-evolving sites removed. The amino acid sites in the 295-marker concatenated matrix were sorted to ten rate categories, with category 10 being the fastest evolving sites. We sequentially removed the fast-evolving sites and perform RAxML analysis with 100 bootstrap replicates and PROTGAMMALG model. The arrangement of (O, (Z, (M, D))) was recovered in all the RAxML analyses. Bootstrap higher than 50 and posterior probabilities higher than 0.5 are shown. D – Dikarya; M – Mucoromycota; O – *Olpidium bornavanus;* Z – Zoopagomycota.

| TIGER rate category removed | None | 10 | 9,10 | 8,9,10 | 7,8,9,10 | 6,7,8,9,10 | 5,6,7,8,9,10 |
|---|---|---|---|---|---|---|---|
| (O, Z, M, D) | 100 | 100 | 100 | 100 | 100 | 100 | 100 |
| (Z, M, D) | 89 | 72 | 82 | 77 | 77 | 70 | 82 |
| (M, D) | 89 | 72 | 85 | 82 | 79 | 74 | 82 |

**Table S4**. Fossil calibrations and gene family GH28 expansion calibration used in MCMCTREE analyses.

| Clade | Fossil | Calibration Age | Calibration Constraint Setting | Ref. |
|---|---|---|---|---|
| Blastocladiomycota | *Palaeoblastocladia milleri* | 407 | lower minimum bound | 1 |
| Chytridiomycota | *Krispiromyces discoides* | 407 | lower minimum bound | 1 |
| Endogonaceae | *Jimwhitea circumtecta* | 247 | lower minimum bound | 1 |
| Mucorales | *Protoascon missouriensis* | 315 | lower minimum bound | 1 |
| Ascomycota | *Paleopyrenomycites devonicus* | 407 | lower minimum bound | 1 |
| Basidiomycota | Clamp connections | 330 | lower minimum bound | 1 |
| Chytridiomycota + *Olpidium* + Zoopagomycota + Mucoromycota + Dikarya | GH28 expansion in Fungi | 1100 | upper maximum bound | 2 |


1) Taylor TN, Krings M, Taylor EL. 2015. Fossil Fungi. Academic Press, London, 382 pp.

2) Chang, Ying et al. 2015. "Phylogenomic Analyses Indicate That Early Fungi Evolved Digesting Cell Walls of Algal Ancestors of Land Plants." *Genome Biology and Evolution* 7(6):1590–1601.
